# Comparative effectiveness of probiotic-based formulations on cecal microbiota modulation in broilers

**DOI:** 10.1101/844613

**Authors:** D.R. Rodrigues, W. Briggs, A. Duff, K. Chasser, R. Murugesan, C. Pender, S. Ramirez, D. Petri, L.R. Bielke

**Affiliations:** Department of Animal Sciences, The Ohio State University, Columbus, Ohio, United States of America; BIOMIN America Inc., Overland Park, Kansas, United States of America; BIOMIN Holding GmbH, Getzersdorf, Austria

## Abstract

The potential of probiotics to manipulate the intestinal microbial ecosystem toward commensal bacteria growth offers great hope for enhancing health and performance in poultry. This study aimed to evaluate the efficacy of five probiotic-based formulations in modulating cecal microbiota in broilers at 21 and 42 days of age. Probiotics investigated included a synbiotic (SYNBIO), a yeast-based probiotic (YEAST), and three single-strain formulations of spore-forming *Bacillus amyloliquefaciens* (SINGLE1), *B. subtilis* (SINGLE2) and *B. licheniformis* (SINGLE3). Metagenomics analyses showed that the cecal microbiota of SINGLE, YEAST, and MULTI supplemented birds had low diversity compared to the control diet without probiotics (CON) at 21d. At the same age, birds fed SYNBIO had a high population of *Lactobacillus salivarus* and *Lachnospiraceae*, which was reflected in a singular microbial community illustrated by beta-diversity analyses. By 42d, there were no differences in alpha or beta-diversity in the microbiota of probiotic-treated broilers compared to CON. Birds treated with SINGLE2 had a lower abundance of lactic acid bacteria, while a higher (*p*<0.05) population of unidentified *Lachnospiraceae* and *Ruminococcaceae*. Although the investigated probiotic formulations shared similar taxonomy, modulation of microbiota was not a core outcome for all probiotic-treated broilers indicating the success of microbiota-based intervention may ultimately be dependent on probiotic mixture and broiler’s age.

## Introduction

The balance between host immune system and intestinal microbial community plays an essential role in health and disease. While pathogen-induced microbiota disruption has been associated with many intestinal and systemic conditions, beneficial bacteria colonization is often linked to high productivity in broilers [1–3]. Therefore, it is becoming increasingly clear that manipulating the microbiota of the gastrointestinal tract (GIT) is an effective strategy to stimulate a healthy balanced microbial community as a means of improving health and performance in poultry [4,5].

In this context, probiotics have been identified as a promising nutritional intervention to promote modulation of GIT microbiota toward commensal bacteria growth [5–8]. However, modification of the microbial population is not considered a general benefit of probiotic supplementation [9]. Mechanisms such as exclusion competition are widespread among probiotic formulations[10]. Nevertheless, effects at the intestinal level are more likely to be strain-specific.

Although several bacterial species and yeasts from *Bacillus, Lactobacillus, Enterococcus, Bifidobacterium, Pediococcus*, and *Saccharomyces* genera have been described as probiotics for broiler chickens [5,11], the probiotic features are more specific to the selected strain than the genus of origin [10]. Additionally, studies with mouse models have shown that taxonomically similar probiotic species produced by different manufacturing methods can exert divergent effects on disease attenuation [12].

Synbiotic supplementation, a combination of both prebiotics and probiotics, has shown potential regarding modification of the gut microbiota and has drawn recent attention [11,13,14]. Since prebiotics are used mostly as a selective medium for the growth of probiotic, the aforementioned modifications in the intestinal ecosystem may occur at the level of individual strains and species [13].

Scientific advances in the field of microbiology have provided crucial insights into the mode of probiotic action. For instance, metagenomics analyses have maximized the knowledge about microbial communities and contributed to developing microbiota-based probiotics. However, there have been inconsistencies concerning the response of probiotic supplementation on the modulation of GIT microbial communities, which underlines the need for a more thorough comprehension of the mechanisms by which probiotics influence the microbiota.

In order to achieve a better understanding of how different probiotic mixtures could affect the GIT microbiota in broiler chickens, this study was conducted to evaluate the efficacy of five probiotic-based formulations in modulating diversity and composition of cecal microbial communities in 21 and 42-day-old broilers.

## Material and methods

### Experimental design and dietary treatments

A total of 720 one-day-old Ross 708 male chicks were allocated to 6 treatments in a completely randomized design. Eight replicates were assigned to each of the treatments with 15 birds per replicate. Treatments were based on supplemental diets including (1) basal diet without probiotics (CON); (2) Synbiotic (0.45 g/Kg ; SYNBIO); (3) Yeast-based probiotic (1.12 g/Kg; YEAST); (4) Single-strain probiotic 1 (0.45 g/Kg; SINGLE1); (5) Single-strain probiotic 2 (0.27 g/Kg; SINGLE2) or (6) Single-strain probiotic 3 (0.45 g/Kg; SINGLE3).

The SYNBIO-based mixture was composed of 2 × 10^11^ CFU/g multi-species probiotic including *Lactobacillus reuteri, Enterococcus faecium, Bifidobacterium animalis, Pediococcus acidilactici*, and a prebiotic (fructooligosaccharide). The formulation YEAST was a non-bacterial probiotic-containing *Saccharomyces cerevisiae* (Moisture 11%, Crude fiber 25%). The single-strain probiotics were composed of spore-forming *Bacillus* spp. Formulation SINGLE1 contained 1.25 × 10^6^ CFU/g of *B. amyloliquefaciens*, while SINGLE2 comprised 10 billion spores/g of *B. subtilis*. Besides, each gram of the SINGLE3 contained 3.20 ×10^9^ CFU of *B. licheniformis*.

Birds were reared from 1 to 42d and housed in floor pens on fresh wood shavings litter with *ad libitum* access to a standard corn-soy diet and water [15]. The feeding program consisted of 3 phases: starter (1-7d), grower (8-21d), and finisher (22-42d). Stater diets were in mash form, whereas the grower and finisher diets were pelleted. All experimental procedures were approved by the Ohio State University’s Institutional Animal Care and Use Committee (IACUC).

### Sample Collection

To investigate the intestinal microbiota composition of probiotic-treated broilers on days 21 and 42, ceca were collected from four birds per pen for DNA extraction and next-generation sequencing (NGS). Cecal contents were weighed and mixed to create pooled samples from two birds (n=16 per treatment for each time collection) for DNA extraction. Next, 0.3g of the mixed digesta was added into a 2.0mL screwcap microcentrifuge tube with 0.2g of zirconia beads (0.1mm). DNA was extracted from each sample, along with pure culture bacterial samples, using the protocol from Arthur et al. [16] with several modifications. After extractions were completed, DNA quality and quantity were measured using a Synergy HTX, Multi-Mode Reader (BioTek, Winooski, VT), and all samples were diluted to a concentration of 20ng/µL for NGS analysis.

### 16S Sequencing Analysis

Generation of PCR amplicon was achieved by amplification of the V4-V5 region of the 16S rRNA gene using 515F and 806R primers (515F: GTGYCAGCMGCCGCGGTAA, 806R: GGACTACHVGGGTWTCTAAT). DNA samples were library prepared for NGS using the Illumina MiSeq platform (2 × 300 bp; Illumina, San Diego, CA, USA) by Ohio State University Molecular and Cellular Imaging Center.

A sequence quality screen was performed to ensure high-quality sequences were submitted to the analysis pipeline. Briefly stated, sequence quality was determined using the FASTQC and MultiQC toolkits. Sequence reads exhibiting a quality score of lower than 20 were removed. Further, low complexity reads, those shorter than 200 bp in length, and mismatched primers were also eliminated. Additionally, reads exhibiting low sequence qualities on either end were trimmed. The pre-processed FASTQ files were then imported to the QIIME2 platform for analysis. The main analytical steps were as follows: firstly, reads were de-multiplexed and classified into their respective samples. Next, additional sequence quality control measures and feature table construction were performed by the DADA2 algorithm implementation in QIIME2.

Quality control measures eliminated reads with barcode errors, along with reads that had more than two nucleotides mismatches, and chimeras. The high-quality sequences originating from the afore-mentioned quality control measures were subsequently clustered together using the q2-feature-classifier plugin with the GreenGenes 13.8 reference database. The resulting feature table was used to calculate diversity metrics and create abundance infographics. Alpha-diversity was measured by Shannon’s diversity index. Beta-diversity was determined by applying pairwise UniFrac distances in QIIME2.

### Statistical Analysis

Kruskal-Wallis pairwise test was assessed to compare differences in the microbial Shannon’s diversity index (H) across treatments. Weighted Unifrac distance metric was used for comparing the cecal beta-diversity between probiotic and non-treated samples (PERMANOVA, *p* ≤0.05). Principal coordinates analysis (PCoA) of unweighted Unifrac distance was addressed to measure the similarity between cecal samples at 21d and 42d. The mean relative abundances of microbial communities were subjected to Analysis of Variance and compared among treatments with a Student’s *t*-test (*p* ≤0.05) using the JMP Pro13 Software (JMP Software, SAS Inc., 2018). For the microbiome plots and heat maps, we used Rstudio software (Version 1.1.463, 2009-2018 RStudio, Inc.).

## Results

Diversity, structure, and composition of cecal microbiome in broilers supplemented with different probiotic formulations were assessed based on 16S rRNA sequencing. A total of 5,348,269 16S rRNA raw sequence reads were obtained. The number of mapped sequence reads of overall samples ranged from 13,545 to 60,125, with a mean of 27,855.82.

### Microbiome diversity and structure

Cecal microbiota alpha-diversity was compared between CON and probiotic-treated groups at 21d (Fig 1A) and 42d (Fig 1B). Shannon’s diversity index indicated that the ceca of SINGLE1, SINGLE2, and YEAST supported a less diverse microbial community comparable with CON at 21d (*p*<0.05). On day 42, although there were no significant differences in the Shannon index compared to CON.

**Fig 1.**
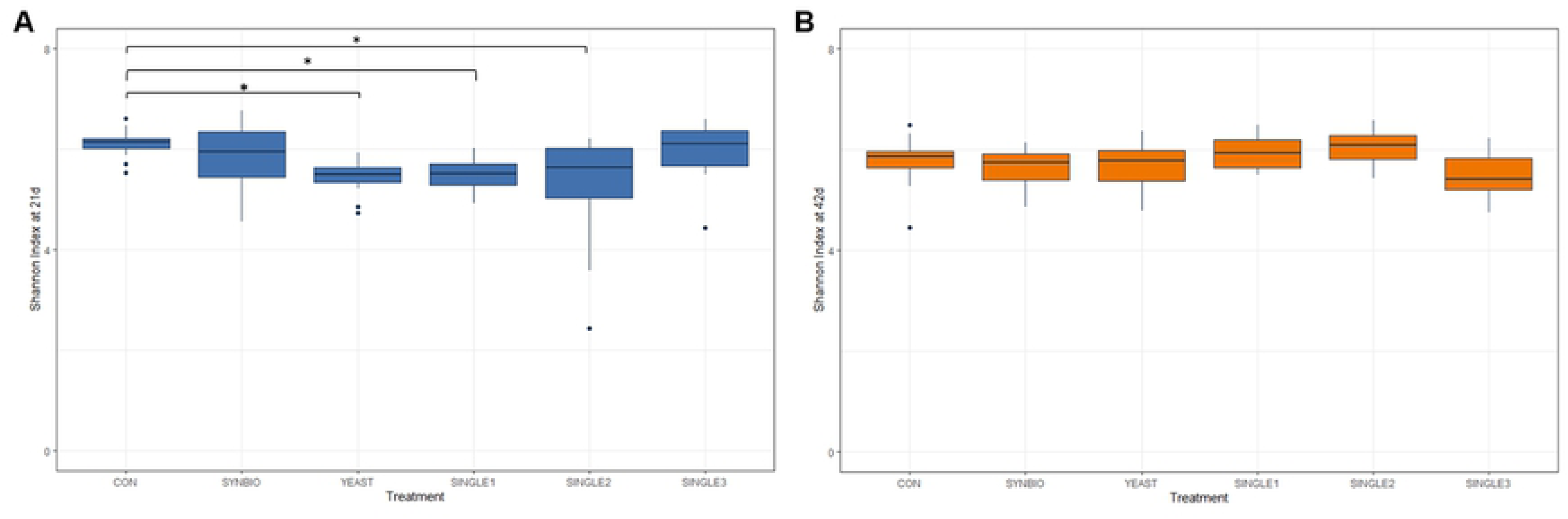
Alpha-diversity in cecal microbiota of broiler chickens treated with different probiotic formulations. (A) Box plots showing the Shannon index of microbial communities sampled in 21 day-old broilers fed basal diet without probiotics (CON), synbiotic (SYNBIO), yeast-based probiotic (YEAST) or single-strain formulations composed of *B. amyloliquefaciens* (SINGLE1), *B. subtilis* (SINGLE2), and *B. licheniformis* (SINGLE3). Panel (B) illustrates the diversity index of microbiota in ceca of 42-days-old broilers.

The cecal microbial community structure of non-treated and probiotic-supplemented groups was investigated at different ages (Figs 2A and B). Weighted pairwise Unifrac distance metric, using the Pseudo-F test, showed significant differences between samples from SYNBIO and CON (*p=*0.02, pseudo-F: 2.62, Permutations: 999) at 21d (Table in S1 Table). Consistently, the hierarchical cluster analysis showed that SYNBIO samples displayed a unique microbial composition with a clear separation from other probiotic treatments by 21d (Fig 3).

**Fig 2.**
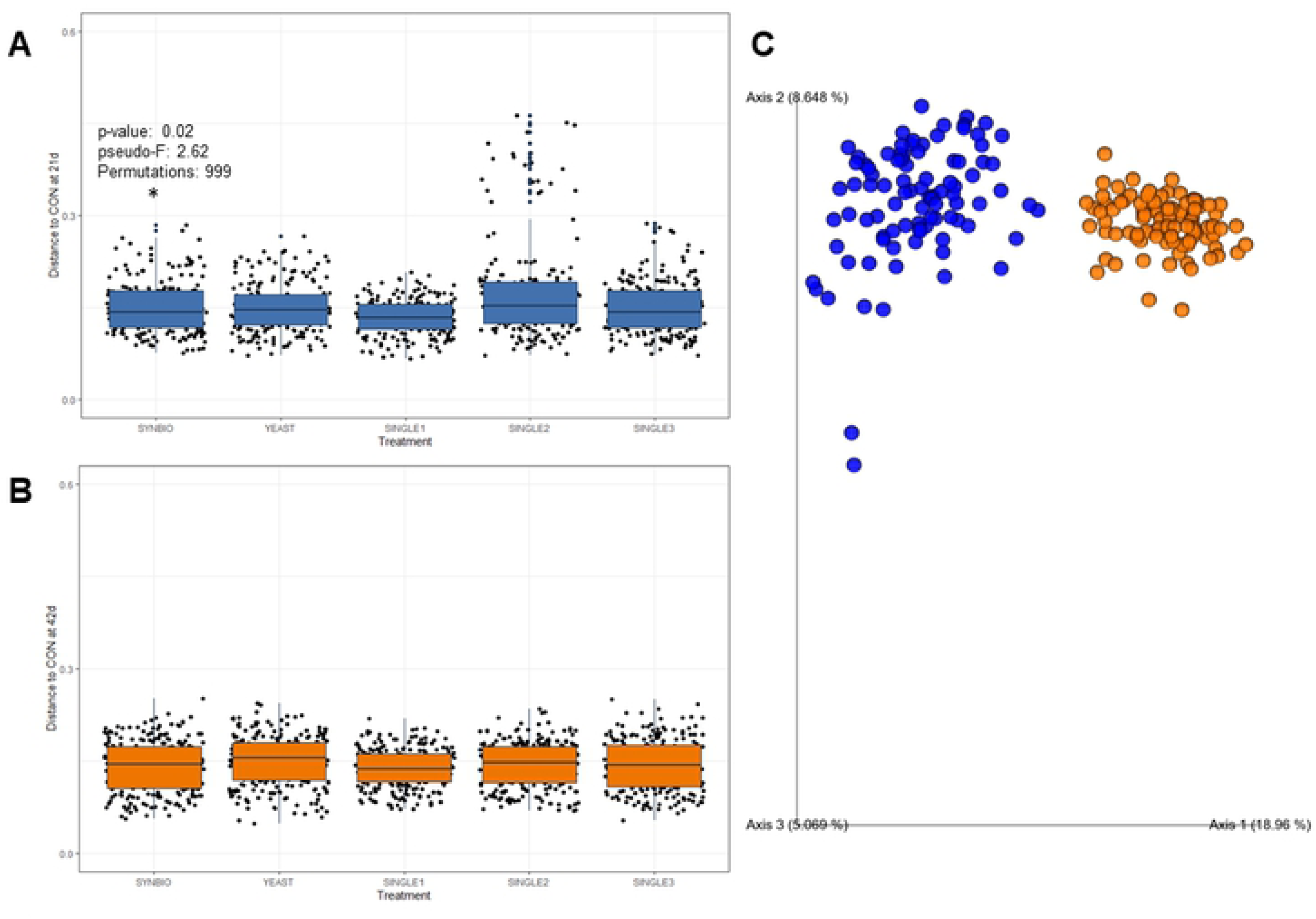
Beta-diversity indices of cecal microbiome in broilers. (A) Pairwise comparison based on weighted Unifrac distances between cecal microbial communities from broilers fed basal diet without probiotics (CON), synbiotic (SYNBIO), yeast-based probiotic (YEAST), or single-strain formulations composed of *B. amyloliquefaciens* (SINGLE1), *B. subtilis* (SINGLE2), and *B. licheniformis* (SINGLE3) by 21 and (B) 42 days of age. Principal coordinate analyses plot derived from unweighted UniFrac confirmed bacterial community differences centered on bird’s age (*p*<0.05, PERMANOVA, blue = 21d, orange = 42d).

**Fig 3.**
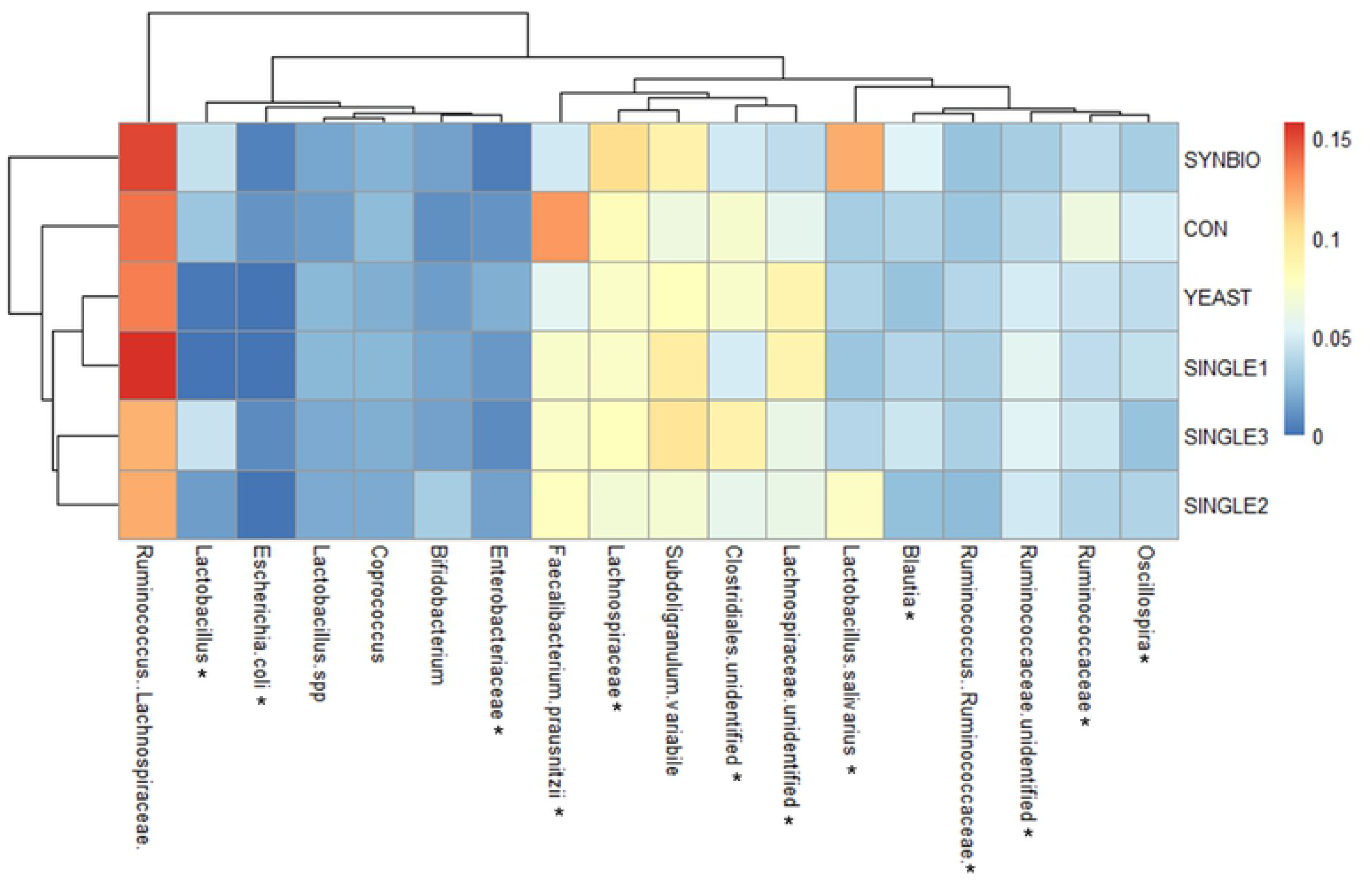
Double hierarchical dendrogram and heat map of cecal bacterial communities from 21-day-old broilers. The taxonomic association of bacterial communities is represented in the columns. Hierarchical clustering in the rows is based on the composition similarity between samples. Statistical differences (*p*<0.05) between groups were reported for each bacterial population (*).

Furthermore, as shown in Fig 2C, there was a clustering of samples based on the birds age. The PCoA plots illustrate the predominant role of age in driving microbiome composition regardless of the probiotic supplementation effect. This dissimilarity between samples from 21 and 42 day-old broilers, revealed by unweighted metric analyses, is further supported by taxonomic-based analyses.

### Microbial community composition

To identify the impact of different probiotic formulations on cecal microbiome makeup, the16S rRNA gene-based microbial profiling was calculated as a percent of mapped reads. As displayed in Figs 3 and 4, the addition of probiotic-based feed resulted in relative abundance changes of key cecal microbial taxa in broilers.

**Fig 4.**
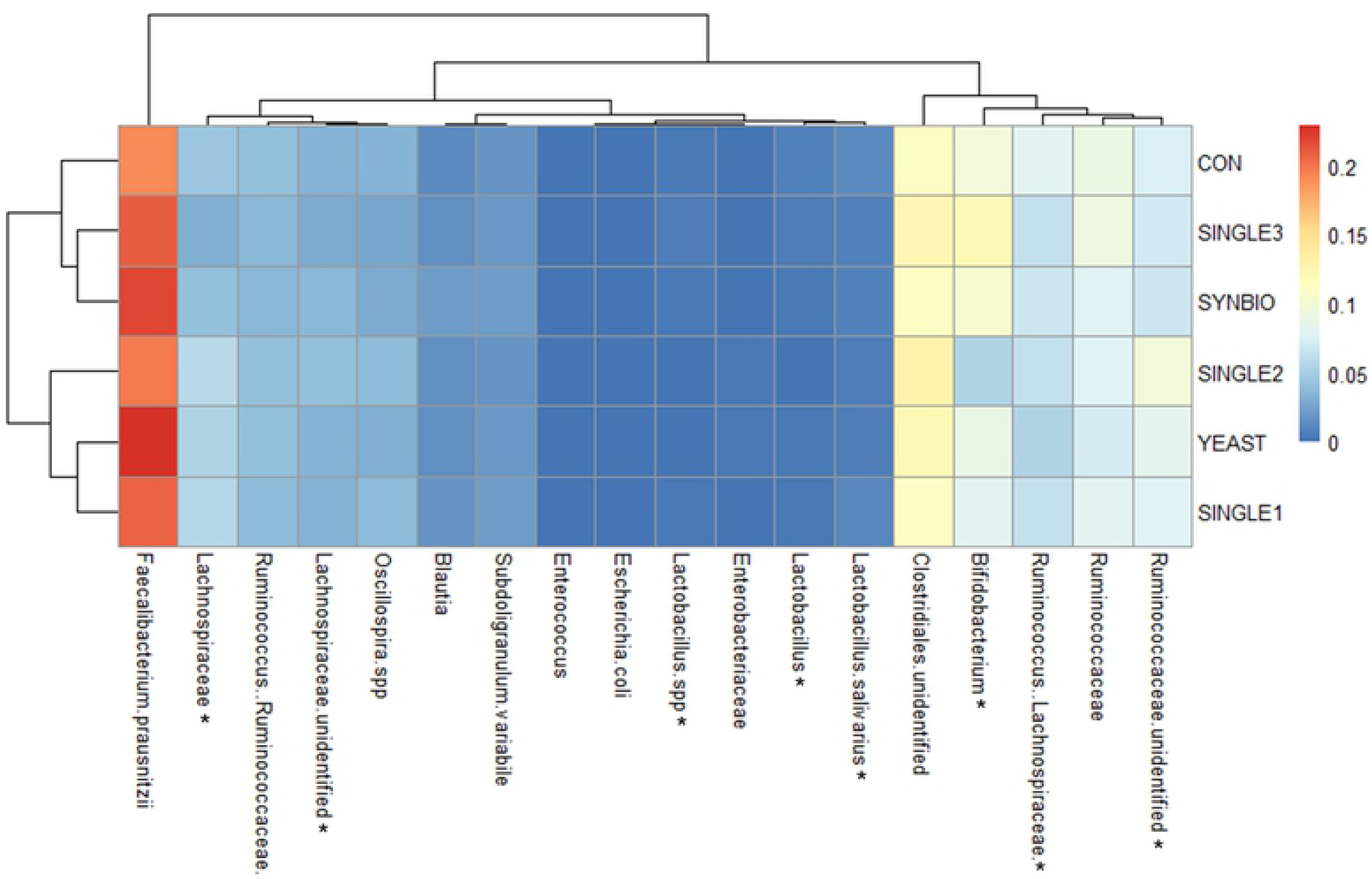
Clustered heat map based upon the predominant bacterial families, genera, and species identified in the ceca microbiome of 42-day-old broilers treated with different probiotic formulations. Statistical differences (*p*<0.05) between groups were reported for each bacterial population (*). Diet without probiotics (CON), supplementation with synbiotic (SYNBIO), yeast-based probiotic (YEAST), and single-strain formulations composed of *B. amyloliquefaciens* (SINGLE1), *B. subtilis* (SINGLE2), and *B. licheniformis* (SINGLE3).

In this study, more than 99.90% of bacterial rDNA sequences were assigned to the domain Bacteria. Dominant bacterial families found in ceca belonged to Clostridiales. Regarding these families, supplementation of SYNBIO increased (*p*<0.05) the population of *Lachnospiraceae*, whereas SINGLE1 and SINGLE2 had significantly reduced *Ruminococcaceae* in ceca relative to CON at 21d (Table in S2 Table). At the same age, *Faecalibacterium prausnitzii* was significantly decreased in SYNBIO and YEAST, while unidentified *Lachnospiraceae* was higher in the microbiota of SINGLE1 and YEAST (*p*<0.05; Fig 3).

Changes in abundance of genus and species from Lactobacillales were also seen between treatments by 21d. The addition of SYNBIO in feed changed the cecal microbiota composition of broilers by increasing (*p*<0.05) the population of *Lactobacillus salivarius*, while YEAST and SINGLE1 reduced (*p*<0.05) the percentage of *Lactobacillus* and *Enterococcus* compared to CON.

Nonetheless, shifts in the relative abundance of species from *Enterobacteriaceae* were observed. Interestingly, although SINGLE1, SINGLE2, and YEAST had the numerically highest population of *Enterobacteriaceae* members, *Escherichia coli* was significantly lower in these treatments compared to CON birds at 21d.

By 42d, there were no broad influences of probiotic supplementation on cecal microbial profile, although a microbial succession pattern was evident (Fig 3 and 4). A considerable increase in *Bifidobacterium* and *Faecalibacterium prausnitzii* populations were detected, but there were no significant shifts across treatments when compared to CON. Notably, the microbiota of SINGLE2-treated birds had a lower abundance of *Lactobacillus, Lactobacillus salivarus*, and *Lachnospiraceae*, while higher (*p*<0.05) population of unidentified *Lachnospiraceae* and *Ruminococcaceae* when compared to CON.

## Discussion

In this study, the impact of different probiotics in modulating the composition and structure of cecal microbial community in 21 and 42-day-old broilers was investigated. Results found here indicate that the effectiveness of probiotics in shaping the cecal microbiota is tied to specific microbial formulation and bird’s age.

We found that the probiotic mixtures tested differently affected richness, evenness, and structure of microbial communities in ceca of broilers at 21d. Supplementation of SINGLE1, SINGLE2, and YEAST reduced the cecal bacterial diversity, whereas SYNBIO and SINGLE3 had similar Shannon index compared to CON. These results disagreed with Wang [17] who reported that supplementation of probiotics in the feed promotes higher biodiversity of intestinal microbiome of poultry. However, as excellently reviewed by Reese and Dunn [18], livestock can have high performance with low-diverse GIT microbiota. This prediction is supported by the notion that the host immune system may limit microbial diversity, given that not all microbes are beneficial [19]. In addition, the overabundance of high-functioning commensal bacteria may lead to a low level of diversity in the intestinal ecosystem [8]. Nevertheless, stress conditions or GIT pathogen colonization can induce a reduction of microbial diversity in poultry [20,21]. The loss in diversity driven by a dysbiotic microbiota is found with a carriage of commensals such as lactobacilli reduced, while the level of *Enterobacteriaceae* increased [22,23]. Here, in this study, there was no pathogen challenge or evident environmental stress imposed on the broilers during the experimental period. Although the cecal *Enterobacteriaceae* population in ceca was not considered high, the abundance of *Enterobacteriaceae* spp. in SINGLE1, SINGLE2, and YEAST was numerically increased at 21d. Even though alpha-diversity is the best predictor to measure richness and evenness of intestinal microbial communities, the available data found in this study is too limited to interpret any substantial conclusion about detrimental implications of the decreasing microbial diversity on health or performance. Addressing this issue would require a deep understanding of diversity-function relationships in host-associated systems [18].

Investigations concerning GIT microbiome diversity and composition have emerged due to evidence that microbiota manipulation may benefit host metabolism, performance, and immune protection to diseases [24–26]. Given that modulation of microbiota can be driven by genetics, diet, environmental conditions, and intestinal pioneer colonization [8,27,28], the age and physiology of organ are identified to play a primary role in influencing composition and diversity of GIT bacterial populations [18,20]. Indeed, ceca have an important function of fermentation and are well known to be the most diverse GIT organ in birds with a predominance of Clostridiales members [29,30]. It has been reported by Lu [29] that the microbial community structure is fairly stable during periods of rapid skeletal growth. For instance, our data showed that there was an age-related difference between 21d to 42d microbiomes, in which microbial shifts were centered on species belonging to *Lactobacillaceae* and *Bifidobacteriaceae*, followed by *Faecalibacterium prausnitzii.*

Probiotics seem to have the greatest effect during the initial development of the microbiota [30]. Of relevance, the supplementation of SYNBIO resulted in a robust modulation of intestinal microbiota with a high population of *Lachnospiraceae* and *Lactobacillus salivarus*, which may explain the differences found in the community structure presented by weighted Unifrac and hierarchical cluster analyses by 21d. However, this uniqueness of the SYNBIO microbial community did not persist through 42d.

As observed in taxonomic analyses, YEAST and SINGLE1 fed broilers had a lower abundance of *Lactobacillus* and *Enterococcus* in the cecal microbiota at 21d. Conversely, by supplementing *Saccharomyces cerevisiae* to broilers diet, Bortoluzzi [31]) found cecal counts of *Lactobacillus* and *Enterococcus* were similar to birds fed a basal diet without yeast-based probiotics. By 42d, SINGLE2 had a major effect in modulating the composition of the bacterial microbiota by decreasing lactic acid bacteria (LAB) populations and increasing unidentified *Lachnospiraceae* and *Ruminococcace*ae. Following this study, Ma et al. (2018) reported that the addition of *Bacillus*-based probiotics affected the cecal microbial composition of broilers by reducing members of *Lactobacillus* along with *Ruminococcaceae*. Unlike other probiotic treatments, in which supplementation resulted in different cecal microbial profiles, the addition of SINGLE3 did not appear to influence the microbiome diversity or composition. Taken together, the results suggest that microbiota modulation was treatment-specific. Also, administration of probiotic-based formulations did not have long-term implications on diversity and composition of resident microbial populations suggesting that young broilers are more responsive to probiotic supplementation than 42-day-old broilers.

While the mode of action of all probiotic strains has not yet been determined, a growing body of evidence suggests that discerning how probiotics modulate intestinal microbiota is essential to define interventional strategies for improving poultry health and performance [5,7]. Also, one of the most promising interests of research and is the concept that microbial additives can be developed to target specific health issues and need [32]. In this context, metagenomics analyses performed in this experiment revealed that the synbiotic supplementation, characterized by SYNBIO, had a better efficiency to modify the microbiota assembly in 21-day-old broilers. A considerable elevation of favorable bacteria such LAB and *Lachnospiraceae* was detected and reflected in a singular microbial community illustrated by beta-diversity analyses. The heightened potential of symbiotic formulations is accredited to mechanisms shared by both probiotics and prebiotics. Teng and Kim [33] have reported that prebiotics from inulin group might stimulate the growth and activity of beneficial bacteria by increasing the concentration of short-chain fatty acids and lactic acid in the ceca of broilers. It is worth highlighting that the intestinal colonization of LAB in poultry has been associated with reduction of pathogens, higher performance, and development of the immune system [8,14,34,35]. Colonization of LAB may also lead to enhanced energy and mineral recovery from nutrients and result in better digestive efficiency [36,37].

In summary, not all probiotic-based formulations tested here had a core benefit on the modulation of microbiota. Dietary supplementation of SYNBIO greatly influenced the growth of favorable bacteria and modified the structure of GIT microbial community in 21-day-old broilers. The success of microbiota manipulation via supplementation of probiotics may ultimately be dependent on specific microbial formulation and birds age. Therefore, prospective studies are warranted to identify associations between specific probiotic formulations and microbiota phenotypes as a means of increasing our understanding of how microbiota-based interventions could offer opportunities to affect health and growth parameters in broiler chickens.

## Supporting information

**S1 Table. Pairwise comparison based on weighted Unifrac distances between cecal microbial communities from broilers**

**S2 Table. Taxonomy of cecal microbial communities in broilers at 21 and 42 days of age**

